# Quantitative Imaging of Receptor-Ligand Engagement in Intact Live Animals

**DOI:** 10.1101/228072

**Authors:** Alena Rudkouskaya, Nattawut Sinsuebphon, Jamie Ward, Kate Tubbesing, Xavier Intes, Margarida Barroso

**Author notes:** Correspondence should be addressed to M.B. or X.I.

## Abstract

Maintaining an intact tumor environment is critical for quantitation of receptor-ligand engagement in a targeted drug development pipeline. However, measuring receptor-ligand engagement *in vivo* and non-invasively in preclinical settings is extremely challenging. We found that quantitation of intracellular receptor-ligand binding can be achieved using whole-body macroscopic lifetime-based Förster Resonance Energy Transfer (FRET) imaging in intact, live animals bearing tumor xenografts. We determined that FRET levels report on ligand binding to transferrin receptors conversely to raw fluorescence intensity. We then established that FRET levels in heterogeneous tumors correlate with intracellular ligand binding but strikingly, not with ubiquitously used *ex vivo* receptor expression assessment. Hence, MFLI-FRET provides a direct measurement of systemic delivery, target availability and intracellular drug delivery in intact animals. Here, we have used MFLI to measure FRET longitudinally in intact animals for the first time. MFLI-FRET is well–suited for guiding the development of targeted drug therapy in heterogeneous intact, live small animals.

---

The most potent drugs are useless if they are not able to engage their targets in living systems. The majority of failed clinical trials are due to the lack of translation of *in vitro* data of drug-target engagement into intact living organisms. Therefore a main issue in drug development is to quantify target engagement in a non-invasive manner in longitudinal preclinical studies in live animals. There are two main approaches to estimate drug binding in animal studies: *ex vivo* invasive methods, e.g. biochemical, radiotracers, flow cytometry, and immunohistochemistry analysis, and measuring binding of the drug to plasma proteins. The latter proved to be a poor indication of drug efficiency (true target engagement) *in vivo*^1^.

Traditional *in vivo* imaging platforms, including PET, lack the ability to discriminate between unbound and receptor-bound macromolecules^2,3^. Especially in the case of cancer pre-clinical models, where the enhanced permeability and retention (EPR) effect, that results from the aberrant leaky microvasculature and inadequate lymphatic drainage of solid tumors, leads to the accumulation of exogenously added protein ligands and antibodies at the tumor region^4^. Therefore, establishing macroscopic colocalization of labeled ligand or antibodies within the tumor region does not guarantee or even indicate actual ligand- or antibody-target engagement in the tumor cells.

Fluorescence lifetime imaging of Forster Resonance Energy Transfer (FLIM FRET) has found numerous powerful applications in the biomedical field^5^. FLIM quantifies FRET occurrence by estimating the reduction of the fluorescence lifetime of the donor fluorophore when in close proximity (2-10nm) to one or more acceptor fluorophores^5,6^. However, to date, FLIM FRET has been confined to microscopic techniques and its translation to the *in vivo* pre-clinical paradigm is just beginning. Herein, we introduce Macroscopic Fluorescent Lifetime Imaging of FRET (MFLI-FRET) as a unique imaging approach in whole-body *in vivo* applications, since MFLI-FRET is capable of discriminating soluble fluorescently labeled ligands from those bound by dimerized receptors. Hence, MFLI-FRET can report molecular events at the macroscopic level in live, intact animals^6,7^.

Since the levels of homodimeric transferrin receptor (TfR) are significantly elevated in cancer cells due to their increased metabolism and proliferation, Tf has been successfully used for molecular imaging and targeted drug delivery^8,9^. Recently, Tf-based imaging has been shown to detect MYC-positive prostate cancer as well as mTORC1 signaling^10–12^. Previously, we established feasibility of measuring FRET signals *in vitro* and *in vivo*, as a result of intracellular near infrared (NIR)-labeled Tf accumulation ^6,7,13,14^.

In this study, we demonstrate the utility of MFLI-FRET as a unique *in vivo* tool to quantify TfR-Tf binding in a longitudinal manner in intact animals. We validated receptor-ligand target engagement in tumor xenogratfs by comparing FRET levels with the results of histological analysis of ligand binding in excised tumors. We show that despite a high heterogeneity of xenografts regarding their size, morphology and receptor density, FRET donor fraction (FD%) directly and robustly reports on Tf delivery into tumor cells. In contrast, we found no correlation between FD% and TfR levels in all xenografts. This finding suggests that when receptor-mediated tumor drug delivery cannot be estimated via *ex vivo* immunohistochemistry, but that drug delivery is highly correlated with FRET imaging.

By providing a direct and quantitative measure of target engagement in live animals in a non-invasive and longitudinal manner, we expect NIR MFLI-FRET to significantly impact the development of targeted therapeutics. NIR MFLI-FRET should find numerous applications in pre-clinical drug delivery as well more generally in the measurement of protein-protein interactions *in vivo* in intact animals.

## RESULTS

### Whole-body imaging of receptor-ligand engagement

The major goal of this study is to demonstrate that MFLI-FRET directly quantifies intracellular tumor delivery of receptor-ligand complexes in live, intact animals. **Fig. 1** explains the important details that distinguish MFLI-FRET from other imaging modalities. MFLI-FRET detects molecular events at nanometer range via the reduction of the donor fluorophore lifetime up**o**n receptor-ligand binding. At the macroscopic level, MFLI-FRET can image the whole animal body and hence the intact tumor environment in a live mouse. Our imaging platform for preclinical imaging applications^15,16^ is based on an innovative time-resolved wide-field illumination strategy and a time-gated ICCD camera to accurately perform fluorescence lifetime sensing on time-gated data sets with optimal photon counts^6,7,13,14^ (**Fig. 1**).

**Figure 1.**
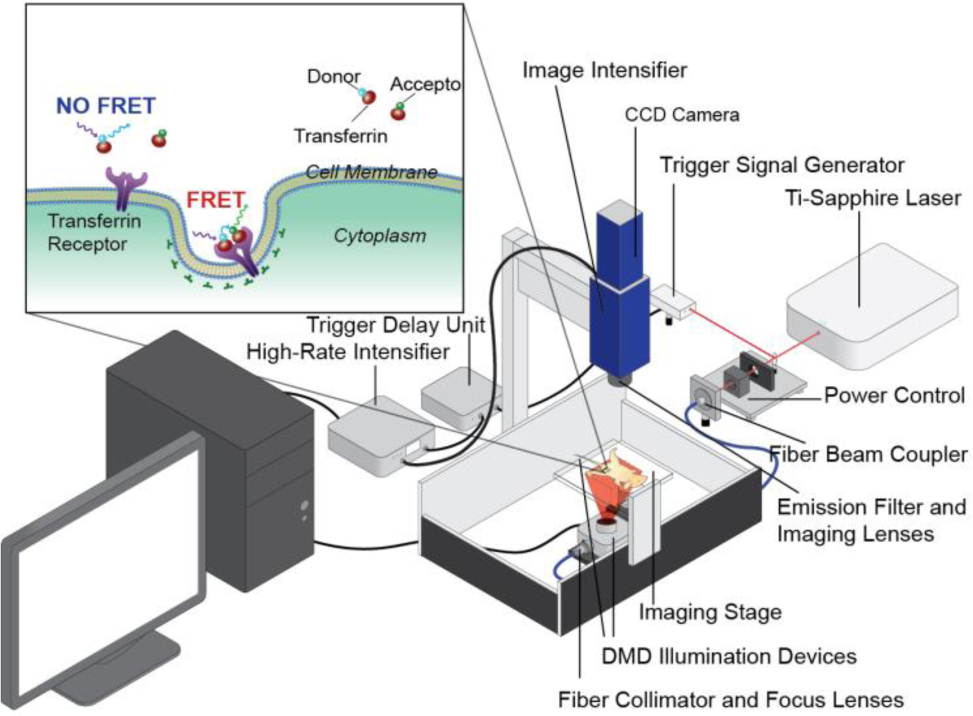
Schematic representation of wide-field macroscopic fluorescence lifetime (MFLI) platform for preclinical application. Inset: Schematic representation of TfR-Tf MFLI-FRET events upon binding of donor-Tf and acceptor-Tf to the homodimeric TfR at the surface of cancer cells. DMD: digital micromirror device; CCD: chargecoupled device.

For this study, we chose the T47D xenograft model since the T47D cancer cell line has a very high level of TfR expression and it has been shown to represent the most common type of breast cancer type in women – luminal A^17^. To demonstrate that MFLI imaging can measure target engagement in a non-invasive manner, three tumor carrying mice (M1-3) were intravenously injected with a mixture of 40 μg Tf-AF700 (donor; D) and 80 μg Tf-AF750 (acceptor; A) and imaged at 24h post-injection (p.i.), which is a standard time point in most optical imaging applications (**Fig. 2**). The data were collected by imaging the whole body of the intact animal, followed by a selection of the tumor and urinary bladder region of interest (ROI) based on the identification of tumor xenografts both by palpation and by fluorescence intensity during the imaging. ROIs were then subjected to lifetime-based data analysis (see Online Methods). Total fluorescently labeled Tf in ROIs (both bound and unbound) are shown as measured by maximum fluorescence intensity (Donor intensity; **Fig. 2a,c**) and FD%, which represents the bound Tf-TfR complex as indicated by reduction in donor lifetime (**Fig. 2b,e**). **Fig. 2d** illustrates the characteristics of the donor fluorescence decay curve of each tumor (T) and urinary bladder (U). Average Tf fluorescence lifetimes in tumors were reduced compared to that in bladders. Very high fluorescent signal in the bladders but minimal FD% were observed consistently (**Fig. 2c,e**). The urinary bladder acts as an internal negative control for each animal, providing a unique gauge non-FRET signal for standardization of FD% levels in tumors. Control staining of excised bladders confirmed that strong Tf fluorescence in the bladders during *in vivo* imaging is associated with detached fluorophore and/or degraded probe in the urine but not with intracellular Tf accumulation (unpublished data). **Fig. 2e** shows that FD% levels in tumors, unlike the donor intensity (**Fig. 2c**), are significantly higher than those detected in bladders indicating TfR engagement of NIR-labeled Tf in tumor cells. Interestingly, pixel distribution clearly shows a bi-modal spatial and quantitative distribution of FD% levels within tumor regions (**Fig 2b,e**). Importantly, our results show FD% levels are not dependent on fluorescence intensity (**Fig. 2**) and FD% estimation remains stable with increased fluorescence intensity (**Supplementary Fig. 1**). These results indicate the ability to infer spatial and intensity level heterogeneity from FD% data distribution in different tumor regions in live animals.

**Figure 2.**
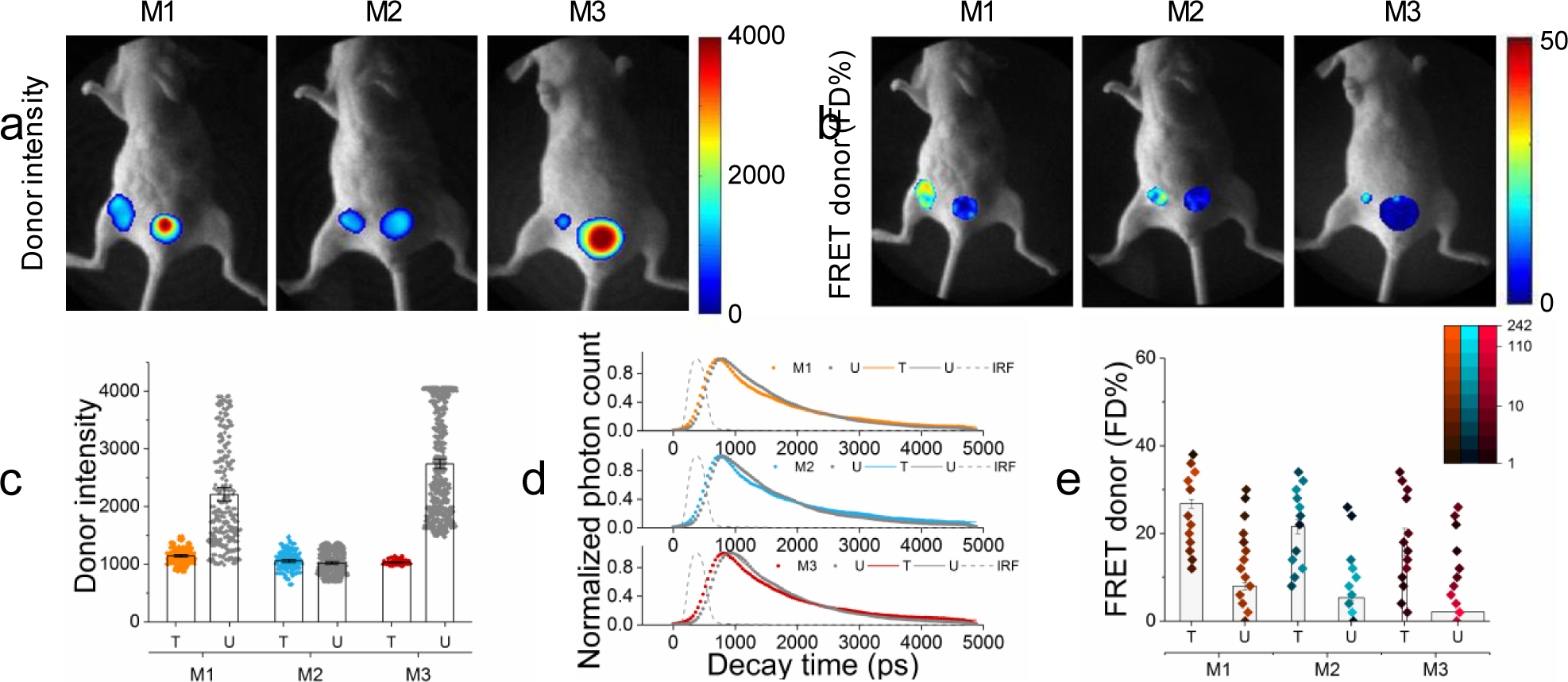
Whole body quantification of ligand-receptor engagement. **(a-b)** Tumor-carrying mice (M1-M3) were injected with 40 μg holo-Tf-AF700 (donor) and 80 μg holo-Tf-AF750 (acceptor) at A:D ratios of 2:1 and subjected to MFLI-FRET imaging at 24h p.i. Panels show Tf donor fluorescence intensity maximum (total Tf, including soluble and bound Tf) **(a)** and FD% levels (bound Tf) **(b)** of live, intact mice using the MFLIFRET imager. Within each mouse image, left ROIs show tumor and right ROIs show bladder fluorescence **(a)** and FD% **(b)** levels. **(c)** Quantification graphs of donor intensity in tumor xenografts at 24h p.i. Data are presented as individual pixels with mean ± 95% confidence interval. **(d)** Donor fluorescence lifetime decay curves of tumors and bladders at 24h p.i. There is a reduction in the fluorescence lifetime in the tumors showing FRET signal compared to the bladders. Bi-exponential fitting is used to retrieve relative populations of donor in the FRETing state (FD%). IRF is the instrument response function. **(e)** Quantification graphs of FD% levels in tumor xenografts at 24h p.i. based on histogram distributions: pseudocolor range indicates pixel intensity frequency using log scale, and boxes represent mean value. The width of boxes indicates the total number of pixels. Error bars represent 95% confidence interval but are too small to display (**Supplementary Table 1**).

### Longitudinal imaging of receptor-ligand engagement

To test the detection sensitivity of *in vivo* MFLI-FRET, we performed longitudinal imaging starting at early p.i. time-points. Four mice were intravenously injected with a mixture of 40μg Tf-AF700 and 80μg Tf-AF750 (M4-6) or 40μg Tf-AF700 alone as a donor only (FRET negative control; M7), imaged and sacrificed at different time points (2h, 6h and 24h p.i.). **Fig. 3a** shows that donor fluorescence intensity of total Tf in tumor ROIs does not necessarily reflect the amount of FD% levels (bound Tf). **Fig. 3b** displays the donor fluorescence decay curves of each tumor at 2h post injection. Importantly, co-injection of donor and acceptor labeled Tf at A:D ~ 2:1 (M4-M6) leads to an increased population of quenched donor fluorophores (**Fig. 3a-b**) and shortening of donor lifetimes (**Fig. 3c**) upon binding of Tf probes to TfR and subsequent internalization into cells. Although FD% quantification at 2h p.i. showed significant variability between xenograft tumors, animals injected with donor and acceptor labeled Tf displayed tumor FD% levels higher than the control animal M7 (**Fig. 3d**). In addition, Tf intensity seems to inversely correlate with FD% indicating quenching of donor upon FRET (**Fig. 3b-d**). Similarly as in the previous dataset (**Fig. 2e**), we observed the bi-modal distribution of FRET FD% signal in tumors (**Fig. 3d**). In agreement, a clear difference is shown between fluorescence decay curves that were determined for representing pixels in each distribution in M6 tumor region at 2h p.i. (**Supplementary Fig. 2**). Moreover, a similar bi-modal distribution was observed when analyzing the MFLI-FRET data with phasor analysis^18^, establishing that such bimodal distribution is not dependent on the data processing methodology. One possible explanation for the FD% bi-modal distribution would be an uneven Tf penetration and binding within the tumor. The non-invasiveness of the MFLI-FRET approach allows for the longitudinal monitoring of target engagement in the same animals as long as fluorescence lasts (**Fig. 3e**). Overall, tumor FRET FD% levels increased dramatically in this experiment during the 2h, 6h and 24 h time course in animals that were injected with both donor and acceptor labeled Tf (A:D 2:1; **Fig. 3e**). In summary, MFLI-FRET imaging is sensitive enough to register FRET signal even as early as 2h p.i. and it can also detect increasing FRET levels over longer periods of time p.i.. Thus, MFLI-FRET imaging should be adequate for a variety of drug-target engagement and drug delivery assessment applications.

**Figure 3.**
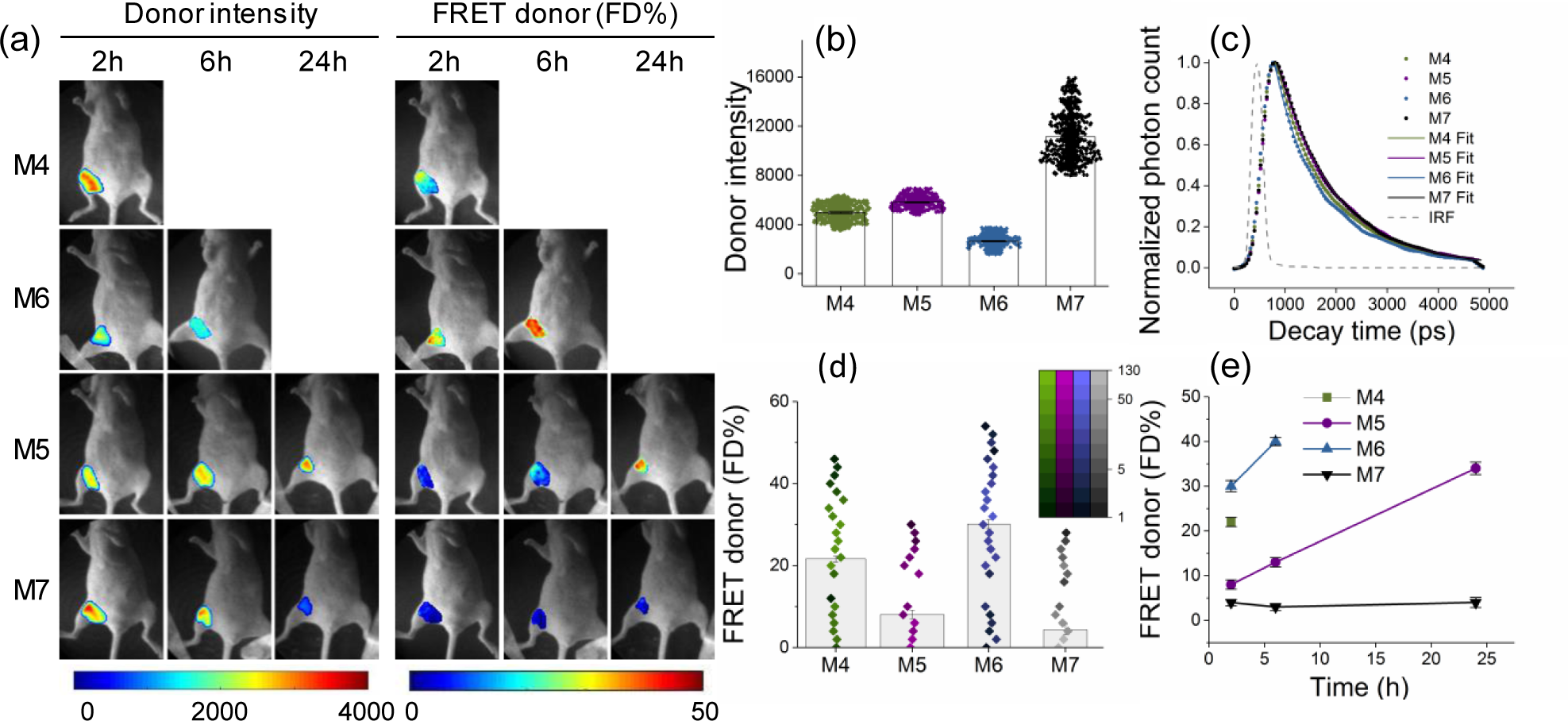
Longitudinal quantitation of ligand-receptor engagement. **(a)** Mice were injected with 40 μg Tf-AF700 and 80 μg Tf-AF750 at A:D ratios of 2:1 (M4-6) and 0:1 (M7) and subjected to *in vivo* wide-field MFLI-FRET imaging at various p.i. time periods. Panels show mice images including tumor ROI with Tf donor fluorescence intensity maximum (total Tf) and FD% levels (bound Tf) of mice using the MFLI-FRET imager as in Figure 2. **(b)** Quantification graph of donor maximum intensity in the tumor regions at 2h p.i., as for Fig. 2c. **(c)** Donor fluorescence lifetime decay curves of tumors at 2h p.i. Bi-exponential fitting is used to retrieve relative population of donor in FRETing state (FD%). IRF is the instrument response function. **(d)** Quantification graph of FD% signal in the tumors at 2h p.i., as in Fig. 2e. 95% confidence intervals are shown in **Supplementary Table 2**. **(e)** Longitudinal imaging of tumors M4-M7 over the course of 24 h MFLI-FRET imaging experiment. Error bars represent 95% confidence intervals.

### Tumor size and morphology heterogeneity

An adequate interpretation of the MFLI-FRET results requires a detailed histological analysis of the tumor xenografts. The excised tumors (M1-M7) revealed a significant heterogeneity of size and morphology (**Supplementary Fig. 3**). Basic morphology of tumors visualized by H&E staining varied from small cancer cell islands in an acellular matrix (tumors M1, M3 and M7), to a reactive lymph node with a massive amount of invading cancer cells (tumors M3 and M4), to densely packed cancer cells in tumors of varying sizes (tumors M2, M5 and M6) as shown in **Supplementary Fig. 3**. Likewise, the xenografts demonstrated a wide range of tumor sizes, with tumors M1 and M2 being at least three times larger than the others (**Supplementary Fig. 4a**). In contrast to much smaller tumors M4 and M6 (**Fig. 3**, **Supplementary Fig. 4a**), FD% in the larger tumors (M1 and M2) (**Fig. 2**) was somewhat lower even at 24h p.i. likely due to penetration issues by NIR-labeled Tf. No correlation between FD% signal and tumor area size across all six tumors was found (**Supplementary Fig. 4b**). Thus, MFLI *in vivo* imaging is fully applicable to a wide range of tumor xenograft sizes and morphologies.

### TfR expression does not correlate with FRET signal

Determining the level of expression of the target receptor is required in a majority of diagnostic assays for anti-cancer therapy selection. Thus, the next question was whether TfR expression level correlates with FD% across heterogeneous tumors. We used the IHC approach to measure receptor expression in paraffin-embedded tumor sections (**Fig. 4a**), since that is the standard approach used currently when assessing total protein expression in tissues. To quantify TfR staining we performed color deconvolution, inversion, and histogram analysis of two separate images per tumor using ImageJ software (**Online Methods**). **Fig. 4b** includes the histogram tumor data, covering the whole tumor area per respective mouse. Since M3 ROI included two small tumor nodules of different morphology shown in **Supplementary Fig. 3** (H&E staining), **Fig 4a** (TfR staining), and **Fig. 5a** (Tf staining), their TfR or Tf integrated densities were combined during analysis. Fluorescence intensity levels per pixel (black lines show mean value of pixel intensity) together with their respective frequency levels were used to calculate the integrated density levels (boxes indicating mean values; **Supplementary Table 3**).The inter-tumoral heterogeneity of TfR expression level and localization was very noticeable (**Fig. 4a**): lymph node M3 aside, the highest TfR level was observed in the largest and most densely packed tumors M2 and M6, where TfR localized mostly to the plasma membrane and cytoplasm. This may be explained by iron deficiency due to oxygen deprivation that occurs in large dense tumors. This hypothesis is supported by *in vitro* studies showing that TfR expression is upregulated by Hypoxia-Induced Factor 1^19,20^. Tumors M1, M4 and M5 displayed somewhat lower TfR levels (**Fig. 4b**). Such a diverse pattern of TfR distribution may be suggestive of a high degree of heterogeneity of Tf penetration inside the tumors. However, when linear regression analysis of FD% at the time of animal sacrifice relative to TfR density levels was performed (**Fig. 4c;** (**Supplementary Table 3 & 5**); no correlation was found between FD% and TfR levels (p=0.32; R^2^=0.2403). Thus, we confirmed the lack of direct relationship between FRET signal (Tf bound to TfR indicating target engagement) and the receptor density levels. This is critical for preclinical studies of drug delivery since patient selection for many types of targeted anti-cancer therapies is based on target receptor overexpression.

**Figure 4.**
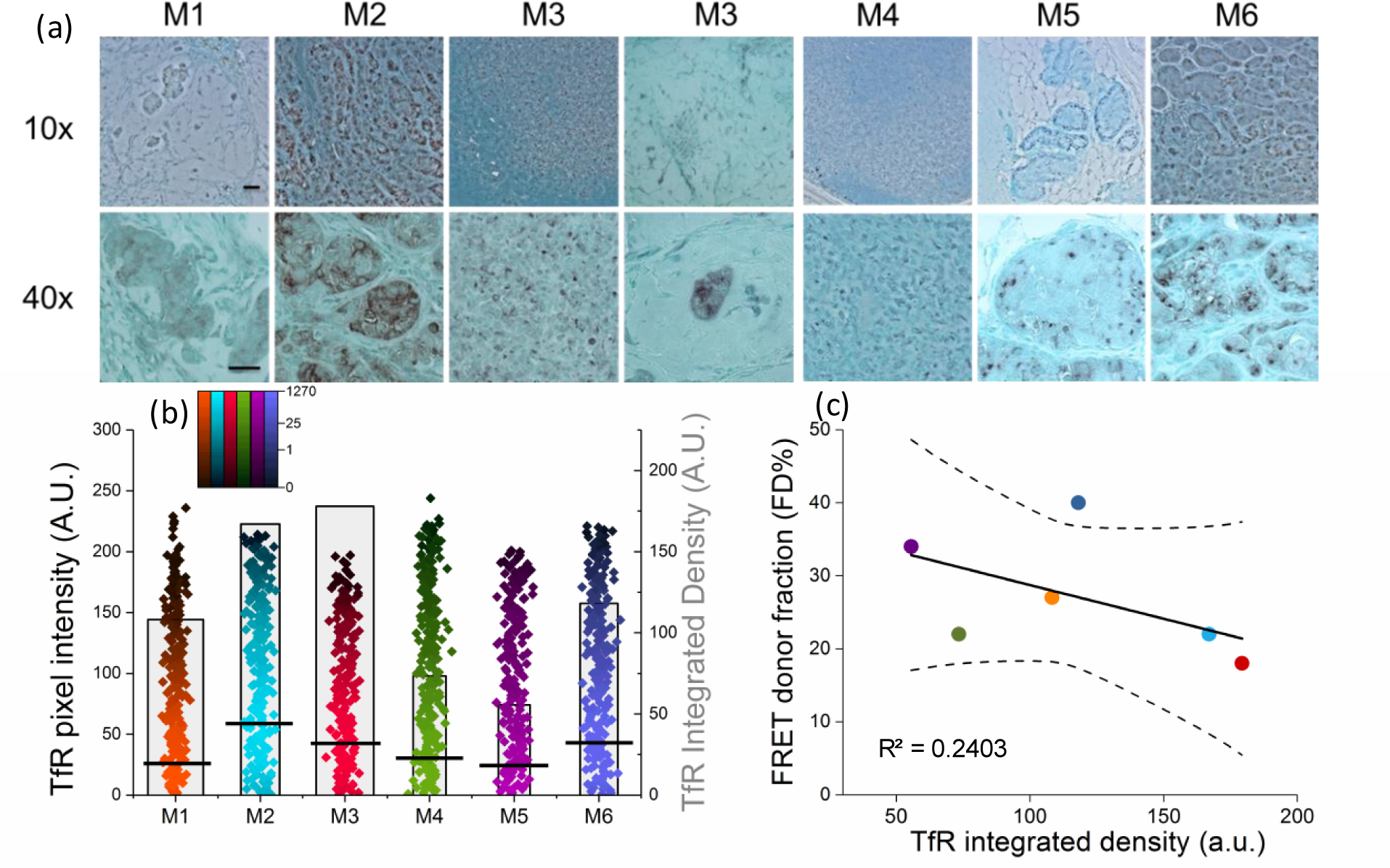
TfR expression level does not correlate with FD% signal. **(a)** IHC staining of TfR in tumor sections using anti-TfR polyclonal antibody, Vector NovaRED as a peroxidase substrate and Methyl Green as a counterstain. The images are shown at 10x (upper panel) and 40x (lower panel) magnification. Scale bar is 100mm. **(b)** Quantification graph of TfR integrated density based on histogram distributions: black lines represent mean values of pixel intensity, pseudo-color range indicates pixel intensity frequency using log scale, and boxes represent mean value of integrated density. Two images per tumor were analyzed, covering the whole tumor. The width of the boxes indicates the total number of pixels. Error bars represent 95% confidence interval but are too small to display (**Supplementary Table 3**). **(c)** Linear regression of the unweighted means of FD% as a function of TfR integrated density. Only FD% values from tumors imaged at the last time point post-injection before sacrifice were included (**Supplementary Table 5**). Regression line and the 95% confidence interval (dotted lines) for the line are plotted. FD% as a function of TfR integrated density levels (slope = -0.08 (SE=0.07) FD%/TfR; p=0.32; intercept=37(SE=9.26); r^2^ = 0.24).

### Tf uptake correlates with FRET signal

The next, most important, step was to assess Tf accumulation within cancer cells via IHC and immunofluorescent staining in relation to corresponding FD% levels. Control experiments were performed to evaluate the contribution of endogenous mouse Tf to the immunofluorescence signal using polyclonal anti-Tf antibody by comparing tumor sections from animals injected with either PBS or human Tf (**Supplementary Fig. 5**). Confocal images collected using the same imaging conditions confirmed that Tf antibody used in this study recognizes primarily human Tf, showing endogenous mouse Tf staining mostly at the background level (**Supplementary Fig. 5**). In this study, both the IHC staining of Tf (**Supplementary Fig. 6**) and the immunofluorescent Tf staining of tumor sections (**Fig. 5a**) confirmed that injected Tf accumulates in cancer and immune cells. In contrast to the trend in TfR levels, the strongest Tf staining was in smaller and less cellular tumors compared to the largest, most densely packed tumor M2. Tumor M6 displayed uneven distribution of Tf within the tissue (**Supplementary Fig. 6** and **Fig. 5a**). Because quantification of IHC Tf staining proved to be ineffective due to the high background of the deconvoluted images (**Supplementary Fig. 6**), whole tissue scans of immunofluorescent Tf staining were quantified and analyzed for integrated density levels (**Fig. 5b**). Again, we compared Tf integrated density levels with corresponding tumor FD% levels of the animal’s last time point imaging prior to sacrifice (**Supplementary Table 4–5**). Remarkably, linear regression analysis using combined data from all six mice showed a strong correlation (p=0.001; R^2^=0.9418). Unlike receptor density measurement, which doesn’t reflect successful ligand binding, MFLI-FRET accurately monitors in real time the receptor-ligand binding and internalization in tumor cells.

**Figure 5.**
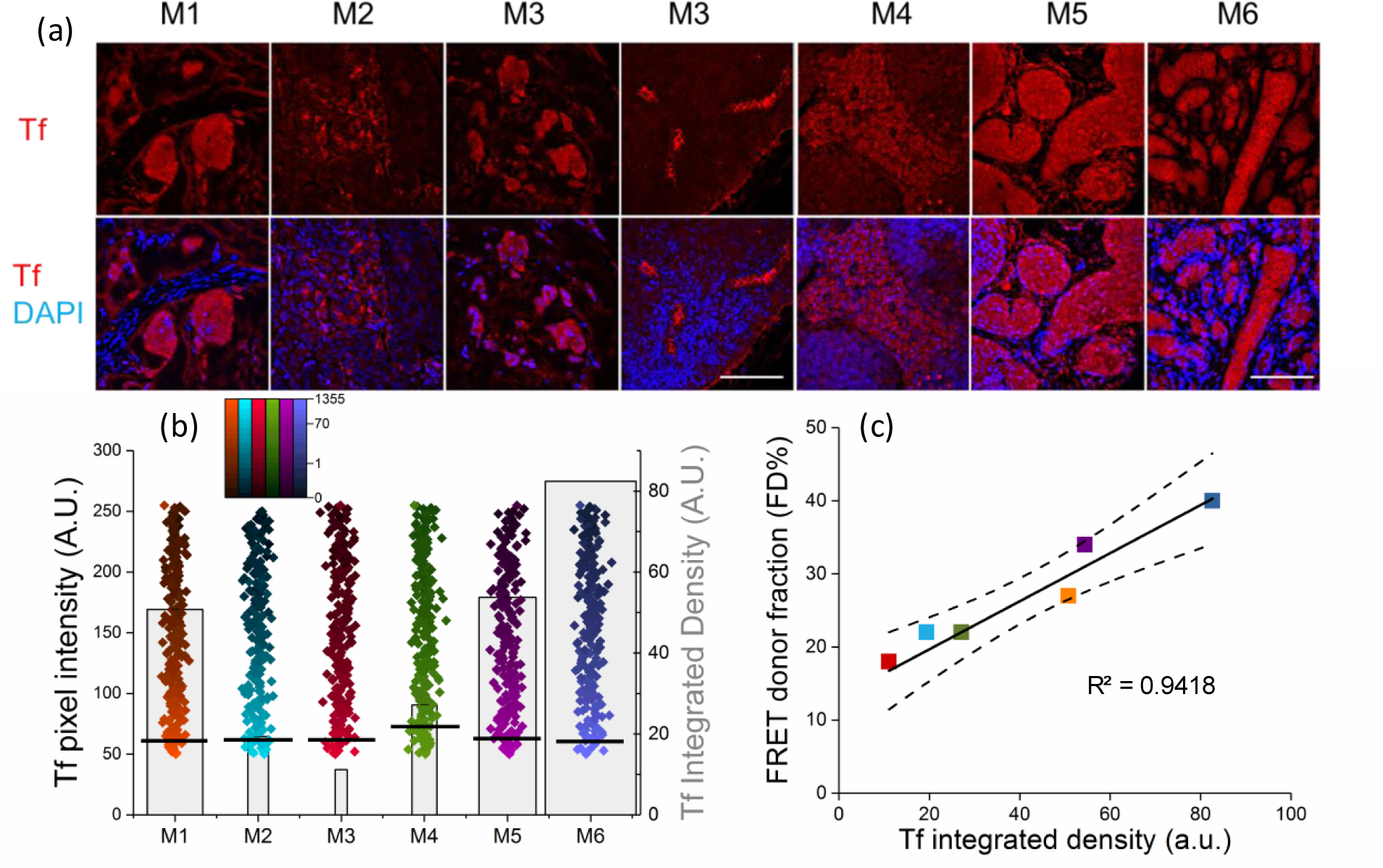
Tf binding to tumor cells correlates with FD% signal. **(a)** Confocal images of Tf immunofluorescent staining in tumor sections M1-M6. Polyclonal anti-Tf antibody was applied to the tissues and visualized with VectaFluor Excel antibody kit. Nuclei are stained with DAPI. Scale bar is 100μm. **(b)** Quantification of whole tissue scans of human Tf immunofluorescent staining was done using Neurolucida system and calculated as integrated density in ImageJ. Black lines represent mean values of pixel intensity, pseudo-color range indicates pixel intensity frequency using log scale, and gray boxes represent mean value of integrated density. The width of the boxes indicates the total number of pixels. Error bars represent 95% confidence interval but are too small to display (**Supplementary Table 4**). **(c)** Linear regression of the unweighted means of FD% as a function of Tf integrated density using data from both imaging experiments. Only FD% values from tumors imaged at the last time point post-injection before sacrifice were included (**Supplementary Table 5**). Regression line and the 95% confidence interval (dotted lines) for the line are plotted. FD% as a function of Tf integrated density levels (slope = 0.3 (SE=0.04) FD%/Tf; p=0.001; intercept=14.75 (SE=1.8); r^2^ = 0.94).

## DISCUSSION

The skyrocketing cost of drug development and high rate of clinical trial failure demand better ways of measuring target engagement *in vivo* to make the drug prioritization process more efficient and reliable^21,22^. We emphasize *in vivo* studies because *in vitro* target engagement measurements are poorly applicable to live organisms, especially to solid tumors, which are notoriously difficult for drug penetration^23^. Traditionally, nuclear medicine has dominated the area of *in vivo* molecular imaging with PET being the main technique to quantitate the targeted drug delivery in live subjects^24^. However, intensity-based PET is limited to the local distribution of the targeted radiotracer and requires multi-compartmental pharmacokinetics models to provide direct information on receptor-ligand interactions^24^. There are a few interesting reports on *in vivo* receptor quantification by dual tracing ^25–27^. Although the dual tracer approach reduces the pharmacokinetic modeling burden, this method still relies on fitting complex models and input of physiological functions despite simplifying assumptions. Successful intravital FLIM-FRET imaging has been used to monitor and quantify drug distribution in tumor cells ^28^. This approach is limited, however, by using cancer cells transfected with Src-FRET biosensor for xenografts, and invasiveness of intravital imaging. A recent study has demonstrated quantification of target engagement of unlabeled drugs using competitive binding with fluorescently labeled companion imaging probes and fluorescence polarization microscopy ^29^. Although this method provides imaging at cellular resolution, it’s limited to intravital imaging and validation of companion imaging probes. In contrast, MFLI-FRET is unique in providing a direct estimate of the fraction of ligand bound to receptor in target cells in intact animals and in a longitudinal manner.

In summary, MFLI-FRET is capable of quantifying target engagement *in vivo* in a simple, elegant and non-invasive and longitudinal manner. One can argue that using fluorescently labeled ligands/ antibodies may reduce their binding abilities, limiting the ability of MFLI-FRET to accurately measure ligand binding in intact animals. However, that would result in underestimation of the target engagement *in vivo*, in contrast to possible overestimation using *in vitro* target engagement assays. Moreover, our results stress the importance of running in parallel *in vivo* and *ex vivo* imaging assays using radioactive or fluorescently labeled ligand for validation purposes.

This study demonstrated a direct correlation between Tf-bound fraction, i.e. FD%, and Tf accumulation in cancer cells as provided by IHC/immunofluorescence imaging, regardless of tumor xenograft heterogeneity in size, morphology, and TfR density. On the other hand, the most striking finding in this work is a general lack of correlation between FD% and TfR expression in the tumors. We speculate that the potential reason for this phenomenon is a tumor microenvironment characteristic for dense solid tumors such as stiff stromal extracellular matrix and EPR effect resulting in elevated interstitial fluid pressure, which together may prevent effective ligand penetration inside the tumor ^23,30,31^. This finding goes against the accepted practice of diagnosis and targeted treatment based on receptor overexpression in tumors. However, in a recent report of Weitsman et al.^32^ the lack of correlation between the overexpression level of HER2 and degree of HER2-HER3 dimerization as an indicator of metastatic likelihood was also shown. It has also been shown that PD-L1 expression does not accurately determine the therapeutic efficacy of anti-PD-1 drugs^33^. This supports our observation that receptor overexpression alone may not be enough for prediction of successful outcome of drug therapy.

The significant advantage of MFLI-FRET imaging is its versatile nature, allowing the use of any ligand/homodimer receptor such as HER2, EGFR, c-MET as well as therapeutic antibodies. In addition, MFLI-FRET has a potential for multiplexing by using simultaneously two target receptors, for example, Tf and anti-HER2 antibody to monitor and quantify drug internalization. Importantly, MFLI-FRET is not limited to breast cancer or other cancer types; it can be applied toward imaging of any disease-targeted drug delivery, such as inflammatory disorders. We recognize though that MFLI FRET is not a universal method for monitoring of target engagement: due to its nature it can be applied only to antibody or ligand/receptor dependent drug delivery. In addition, for dimerized receptors, MFLI-FRET has an inherent limitation to quantify only a fraction of ligand/receptor interaction, up to 50%. Receptor clustering has the opposite effect possibly allowing FRET signal to increase.

This study demonstrates for the first time NIR MFLI-FRET as a robust and direct measure of target engagement by direct assessment of intracellular ligand accumulation in heterogeneous breast cancer xenografts. The results also point out that a high level of receptor expression may not guarantee the efficacy of drug delivery. This unique non-invasive imaging methodology has a potential to greatly benefit the field of targeted drug delivery and to become the gold standard of target engagement quantification.

Furthermore, MFLI-FRET should be as broadly applicable to *in vivo* imaging, as FLIM FRET to *in vitro* fluorescence microscopy. FRET microscopy has been one of the most extensively used imaging techniques in biomedical research, and here we use MFLI methodology to extend NIR FRET to in vivo imaging of small animals.

## METHODS

### Ligand labeling

Human holo Tf (Sigma cat# T8158) was conjugated to Alexa Fluor 700 or Alexa Fluor 750 (Life Technologies cat#A20010 and A20011) according to manufacturer instructions. The degree of labeling of the probes was assessed by spectra analysis using FlexStation 3 (Molecular Devices, Sunnyvale, CA, USA). The average degree of labeling was 2 fluorophores per Tf molecule.

### Cell lines

The human breast cancer cell line T47D (HTB-133) was purchased from ATCC (Manassas, VA, USA) in 2013 and used within one year since the purchase. Cells were cultured in Dulbecco’s modified Eagle’s medium (Life Technologies, cat#11965) supplemented with 10% fetal calf serum (ATCC, 30-2020), 4 mM L-glutamine (Life Technologies), 10 mM HEPES (Sigma) and Penicillin/Streptomycin (50Units/mL/50ug/mL, Life Technologies) at 37oC and 5% CO2. Cells were cultured no longer than passage 20 after thawing. Mycoplasma test was performed once in three months using Universal Mycoplasma Detection kit (ATCC).

### Tumor Xenografts

Tumor xenografts were generated by injecting 10×10^6^ T47D cells in phosphate-buffered saline (PBS) mixed 1:1 with Cultrex BME (Trevigen, Gaithersburg, MD, USA, cat#3632-010-02) into the right inguinal mammary fat pad of female 5-week old athymic nude mice (Crl:NU(NCr)-Foxn1nu, Charles River Laboratories, Wilmington, MA, USA). The use of estradiol pellets to grow T47D tumor xenografts was omitted with the goal of achieving a diverse group of tumor xenografts. The tumors were allowed to grow for 3-4 weeks and were monitored daily. All animal procedures were conducted with the approval of the Institutional Animal Care and Use Committee at both Albany Medical College and Rensselaer Polytechnic Institute. Animal facilities of both institutions have been accredited by the American Association for Accreditation for Laboratory Animals Care International.

### Wide-field Macroscopic Fluorescence Lifetime Imaging Platform for *in vivo* Application

MFLI was performed on a wide-field time-domain fluorescence lifetime imaging tomographic system, as detailed in ^7,16^ (**Fig. 1**). Briefly, the system excitation source was a tunable Ti-Sapphire laser (Mai Tai HP, Spectra-Physics, CA). The spectral range was 690 – 1040 nm with 100-fs pulses at 80 MHz. The laser was coupled to a digital micro-mirror device (DLi 4110, Texas Instruments, TX) which produced a wide-field illumination over an 8×6cm area at 1024 × 768 pixel resolution and with 256 grey scale levels encoding at each pixel. The wide-field illumination was spatially modulated by controlling the DMD via Microsoft PowerPoint to ensure optimal signal detection over the whole animal body^13,14,34^. The detection system was an intensified gated ICCD camera (Picostar HR, Lavision GmbH, Germany). The gate width on the ICCD camera was set to 300 ps. The Instrument Response Function (IRF) and fluorescence signals were collected with 40 ps time interval over a 2 ns time window and a 4.6 ns time window, respectively. The total number of gates acquired was 121 and the maximum range of detection was 4096 photons per pixel per gate. The multichannel plate gain (MCP) employed for signal amplification was set to 550 V for the whole imaging session. In this study, imaging was performed in transmission mode. The laser excitation was set at 695 nm and the emission filter was 720±6.5 nm (FF01-720/13-25, Semrock, IL). The IRF was measured by using a full field pattern projected on diffuse white paper and acquiring the temporal point spread function (TPSF) without using an emission filter. The imaging platform was equipped with an isoflurane anesthesia machine, a warming device, a physiological monitor and a euthanasia system, described in detail in ^7^.

### Fluorescence Lifetime Data Processing

The imaging system generated 3D date cubes (x,y,t; 172×128×121), of which each pixel (x,y) represented fluorescence lifetime decay. The analysis of the fluorescence decay was performed in MATLAB with bi-exponential fitting model: 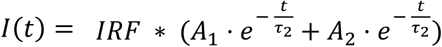, where I was fluorescence intensity at time delay t. τ1 and τ2 were short and long lifetimes of donor fluorophores in FRETing and non-FRETing state respectively. A1 and A2 were relative amplitude of short and long lifetime components. The short lifetime was set to 280±40 ps, and the long lifetime was set to 1050±50 ps which is the estimated lifetime of Alexa Fluor 700 under non-FRETing conditions. The estimation of A1 and A2 in this study was performed using tail-fitting at 99% to 1% of the late time gates. A1 was used to represent FRETing donor fraction (FD%), which was a quantitative parameter for assessment of FRET in vivo. As bi-exponential parameter estimation is well-known to be sensitive to noise, the region of interest used for analysis was defined by a local fluorescence intensity thresholding (any pixel with maximum gate counts>500). Such thresholding is performed to remove any pixel with low-intensity that cannot be reliably used for dual-exponential fitting ^35,36^.

### Immunohistochemistry

Upon imaging, tumors were surgically removed, fixed in 4% paraformaldehyde, paraffin embedded and processed for immunohistochemistry (IHC). Epitope retrieval was performed by boiling deparaffinized and rehydrated 5μm sections in 10 mM Sodium citrate pH 6.0 for 20 min. IHC staining was carried out using a standard protocol from Vectastain ABC Elite kit (Vector Labs, Burlingame, CA cat#PK-6101). Vector NovaRED (Vector Labs) was used as a peroxidase substrate. IHC stained tissue sections were counterstained with Methyl Green (Sigma, cat# M8884). For immunofluorescent Tf staining epitope retrieval was done by boiling tissue sections in 10 mM TRIS pH 9 for 30 min followed by a manufacturer protocol (VectaFluor Excel DyLight 594 antibody kit, Vector Labs, cat# DK-1594). Immunofluorescently labeled tissues were counterstained with DAPI (Invitrogen). Hematoxylin Eosin stain was used for basic histology. All tumor sections were evaluated in a blinded manner by an independent experienced pathologist. Primary antibodies were as followed: rabbit polyclonal Tf 1:2,000 for IHC and 1:200 for IF (Abcam, cat#1223), rabbit polyclonal TfR 1:250 (Abcam, cat# 84036), rabbit polyclonal ER 1:100 (NeoMarkers cat#RB-1493). Brightfield images were acquired using Olympus BX40 microscope equipped with Infinity 3 camera (Lumenera Inc., Ottawa, ON, Canada). Color deconvolution for histograms was performed using ImageJ software. Immunofluorescent images were acquired using Zeiss LSM510 two-photon confocal microscope. For quantification analysis, images of whole tumors tissues were acquired using Neurolucida 64 bit 3D reconstruction and stereology system (MBF Bioscience) and analyzed in ImageJ using a threshold of 50 pixels.

### Statistical Analysis

The statistical significance of the data was tested with unpaired Student’s t tests. Differences were considered significant if the p-value was less than 0.05. Error bars indicate standard deviation. Linear regression analysis and 95% confidence interval were done using OriginPro 2016 software (Northampton, MA).

## Acknowledgements

The authors thank Dr. Stewart Sell (Wadsworth Institute) for xenografts histopathology evaluation, Dr. Paul Feustel (AMC) for the help with regression analysis and Dr. Joe Mazurkiewicz (AMC) for his help with Neurolucida image analysis software. We thank the AMC imaging core facility for the use of the Zeiss LSM510 confocal microscope. We thank the members of the Barroso and Intes labs for their helpful discussions and comments. We also thank Dr. Lamar (AMC) for help with tumor biology. and manuscript discussion. The study was funded by NIH grants R01EB019443 and R01CA207725.

## Author contributions

X.I. and M.B. conceived the original idea. N.S. contributed to the platform instrumentation, optical imaging data collection and analysis and animal study protocol and experiments. A.R. contributed to the animal study protocol and experiments, mice tumor xenograft generation and characterization, optical imaging data collection and analysis, microscopy image collection and analysis and tissue section preparation. K.T. contributed to animal study protocol and experiments and microscopy data collection and analysis. J.W. contributed to image data analysis. All authors contributed the experimental result analysis and manuscript writing.

## Additional Information

Correspondence and requests for materials should be addressed to M.B and X.I.

## SUPPLEMENTARY INFORMATION

**Supplementary Figure 1.**
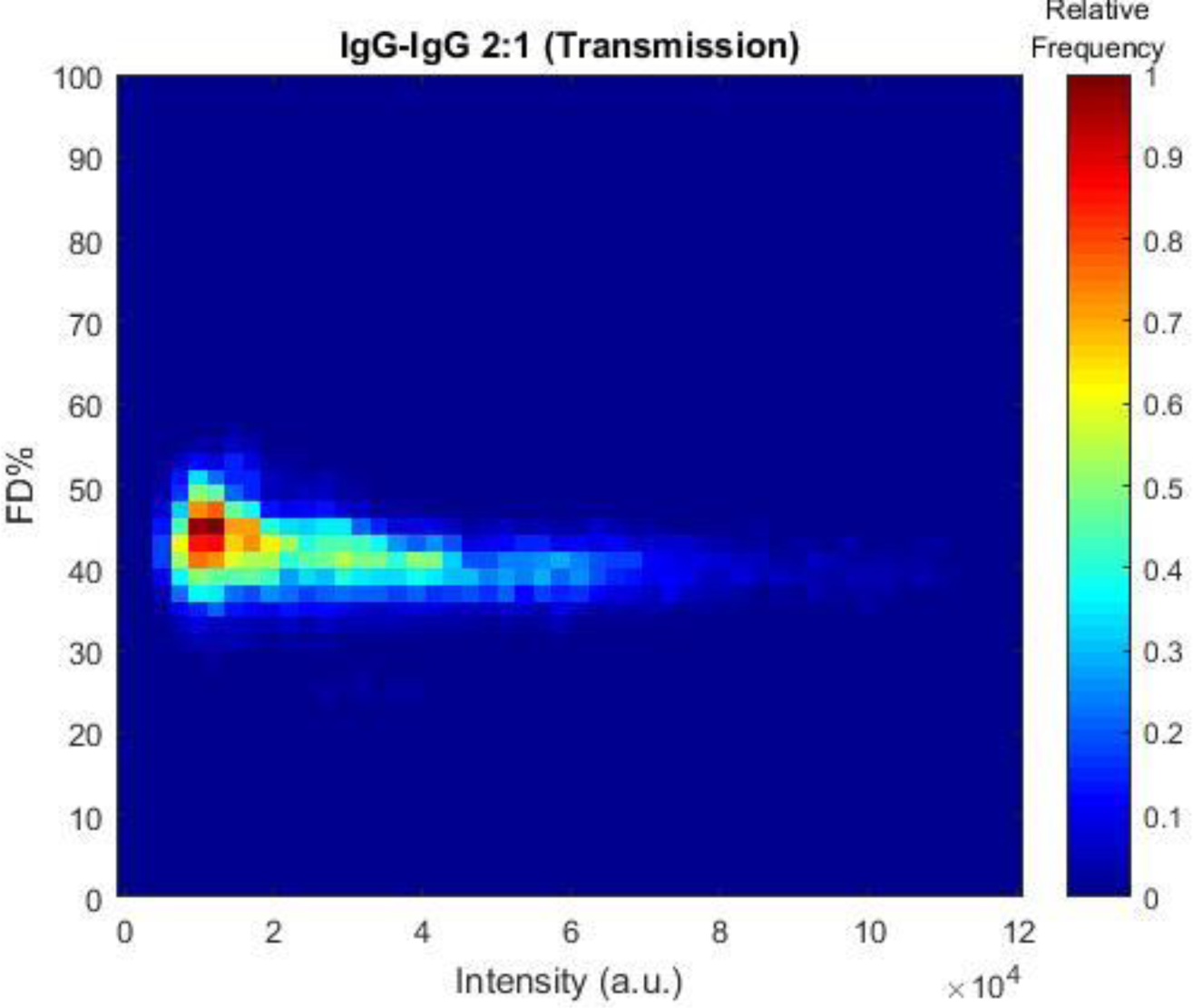
3D histogram (scattered plot + histogram) showing a distribution of FD% and intensity for IgG-IgG binding at A:D 2:1 using transmission MFLI FRET. The total pixel is 18,709.

**Supplementary Figure 2.**
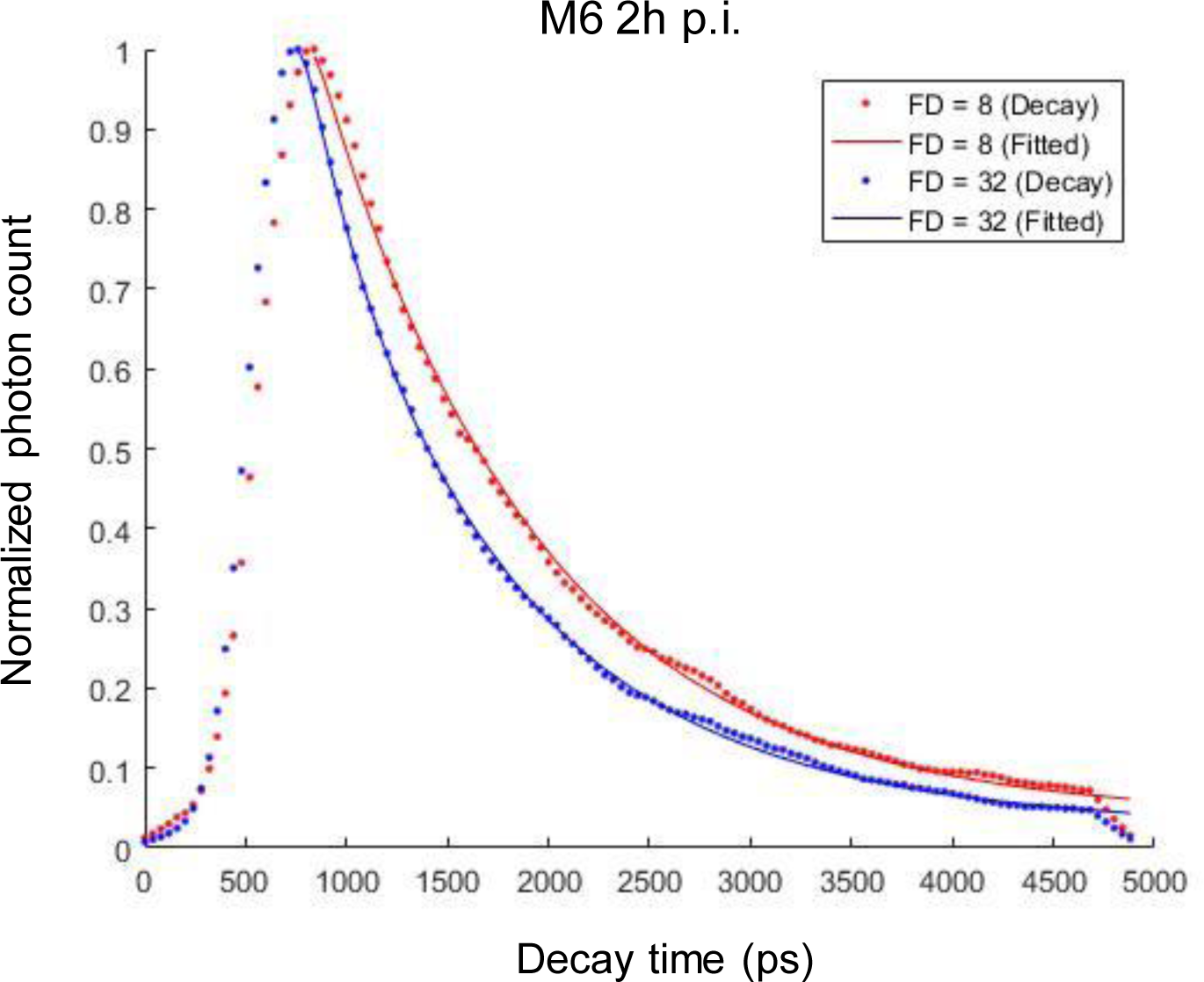
Fluorescence decay (dotted lines) and fitted (solid lines) curves of 2 pixels of M6 at 2h p.i. with different FD% (8%-red- and 32%-blue-).

**Supplementary Figure 3.**
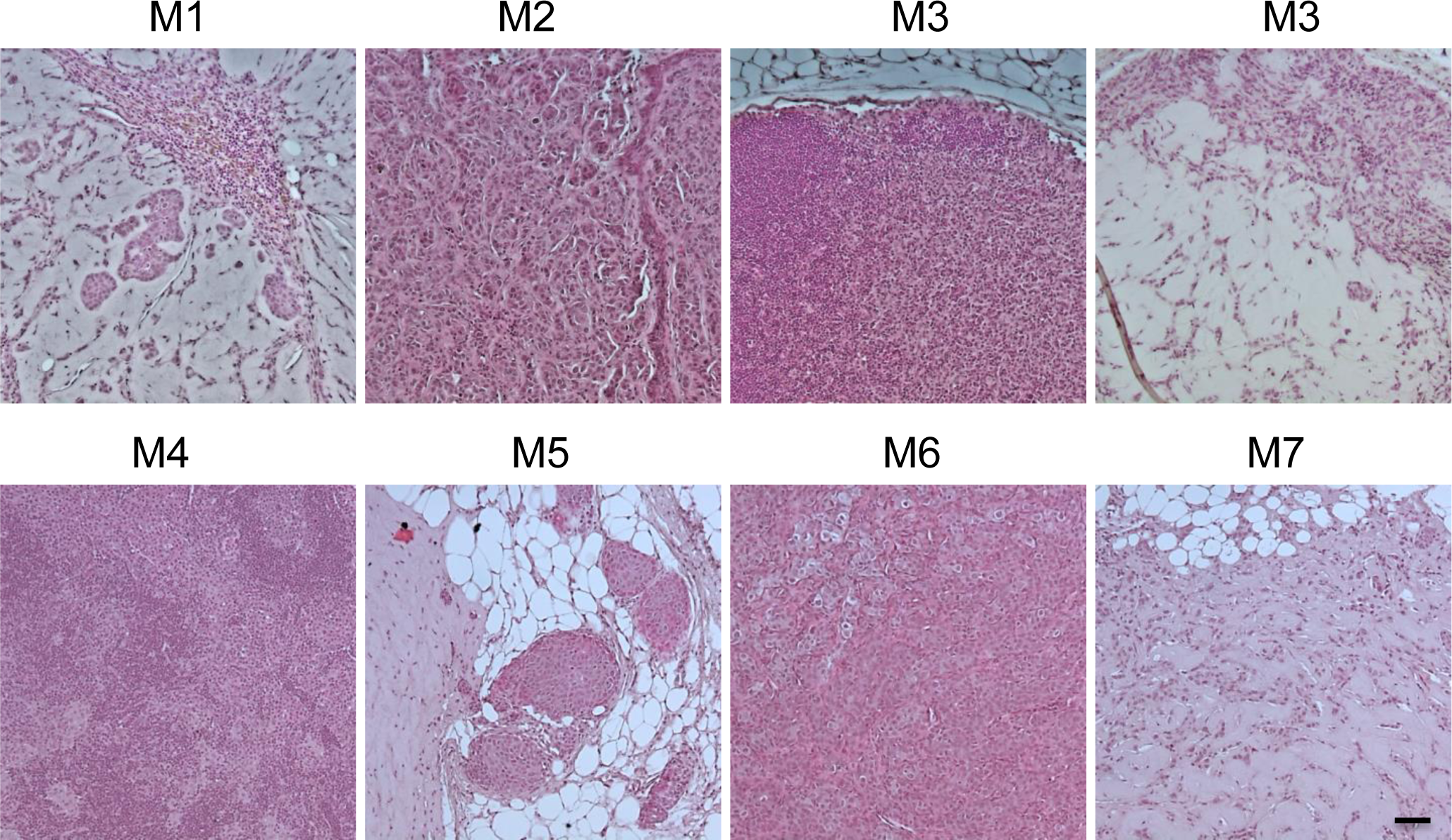
Hematoxylin and Eosin staining for basic morphology of tumors. Scale bar is 100 μm

**Supplementary Figure 4.**
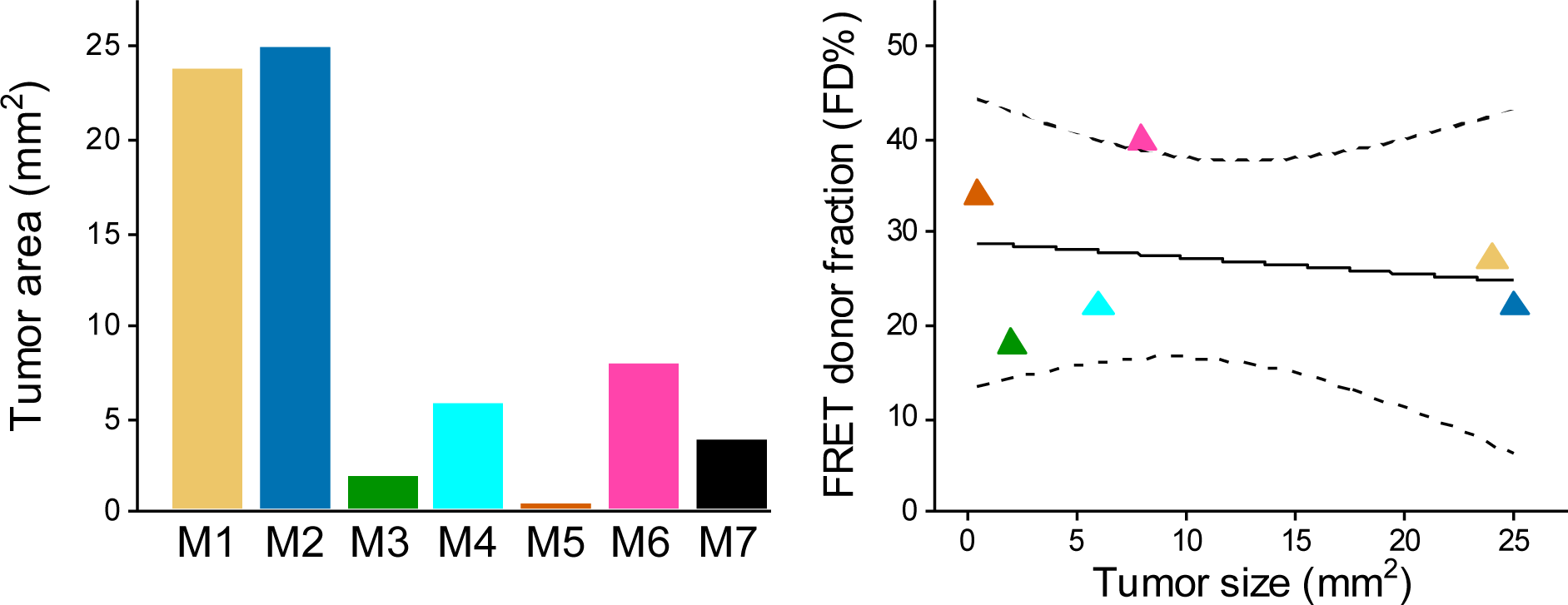
**(a)** Quantification graph of tumor area was determined by measuring approximate dimensions of tumor sections using stage micrometer. **(b**) Linear regression of the unweighted means of FD% as a function of tumor area, slope = -0.13 (SE=0.38) FD%/tumor area; p=0.75; intercept=29 (SE=5.6); r^2^ = 0.028). Regression lines and the 95% confidence interval (dotted lines) for the line are plotted.

**Supplementary Figure 5.**
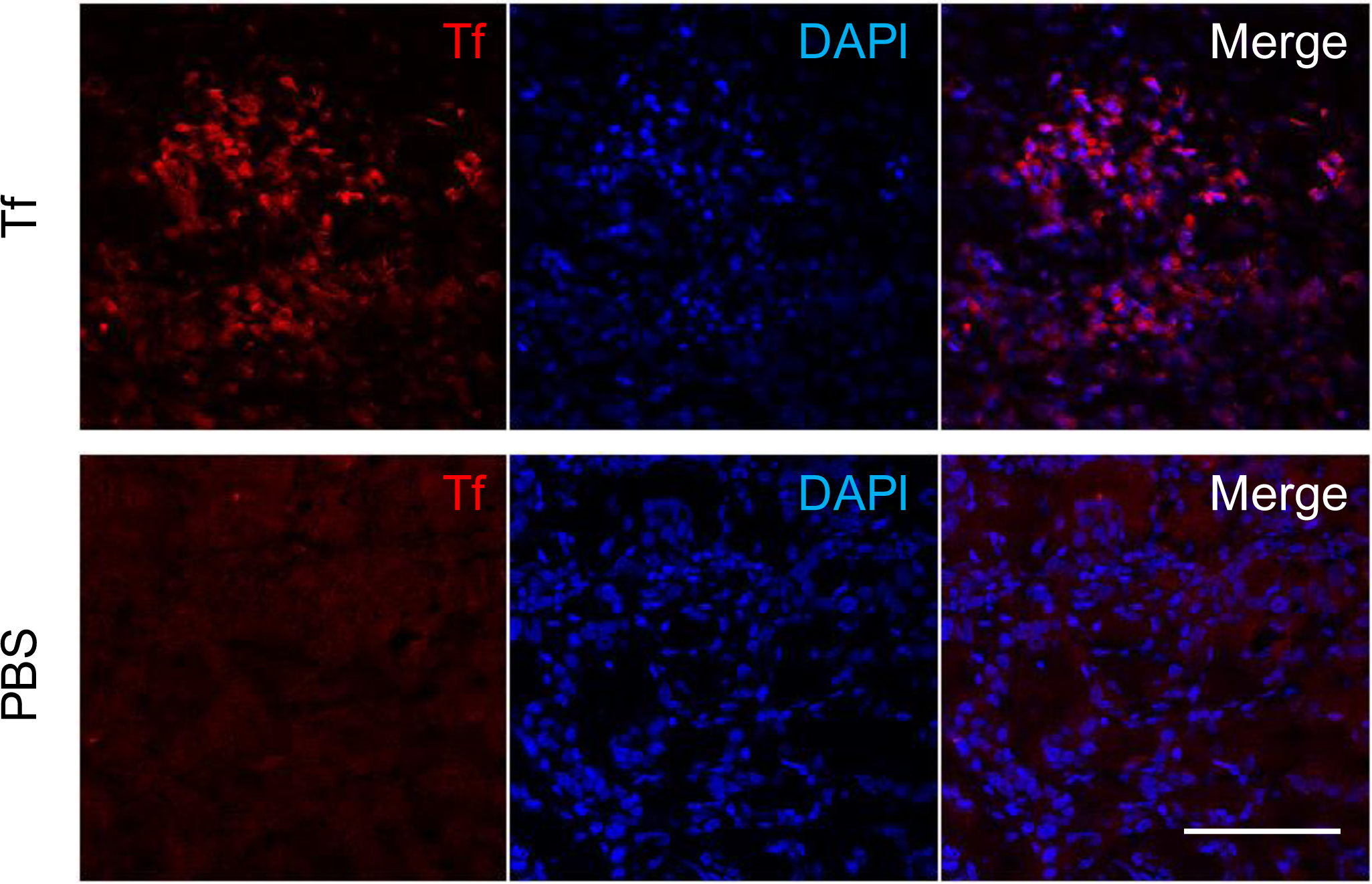
Tf antibody used in this study recognizes primarily human Tf. Confocal images of Tf immunofluorescent staining of T47D tumor xenografts. Nude mice carrying T47D tumor xenografts were tail vein injected with either human Tf (Top panel) or PBS buffer (Bottom panel). Nuclei are stained with DAPI. Scale bar is 100 μm.

**Supplementary Figure 6.**
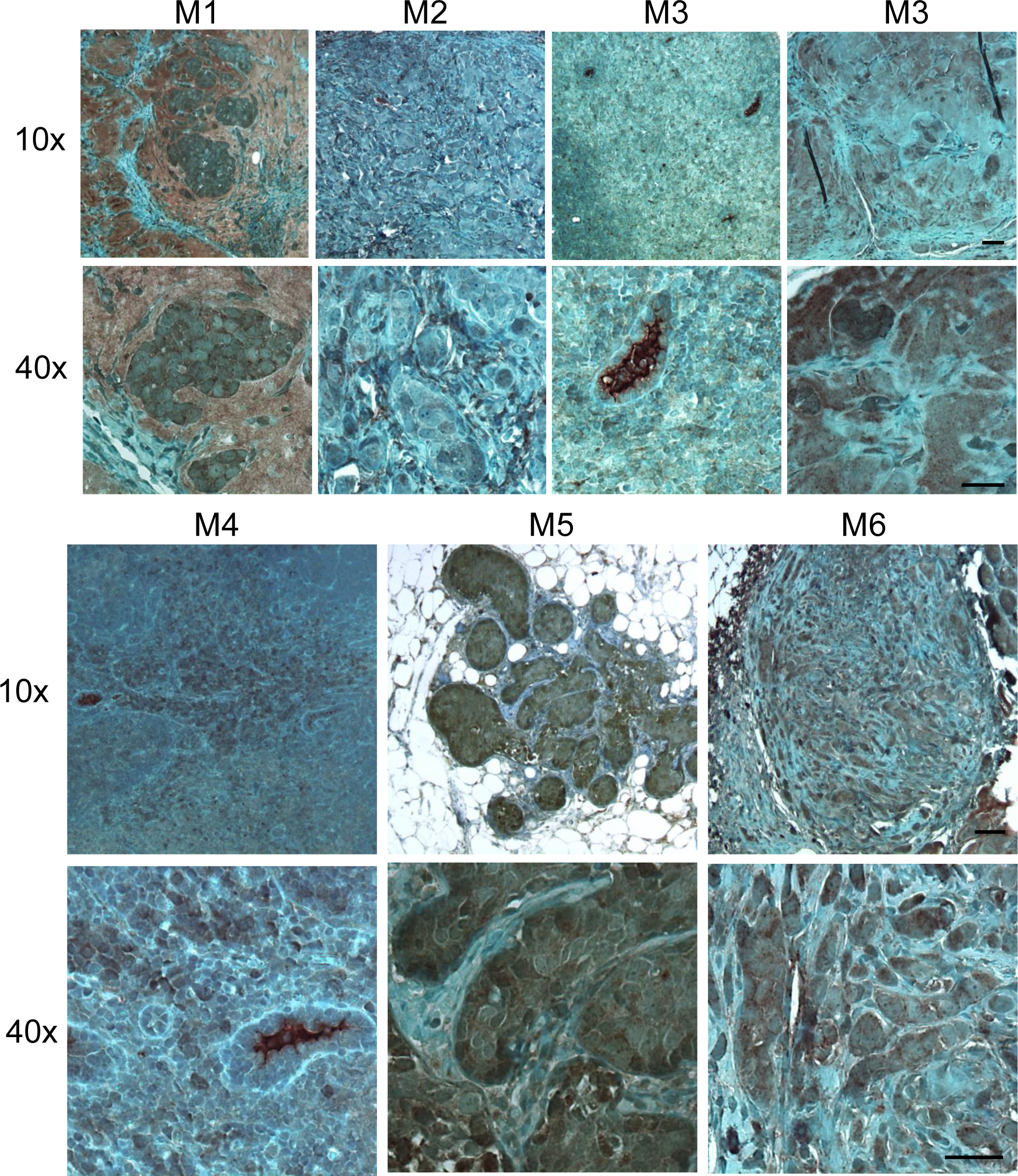
IHC staining of accumulated Tf in tumor sections from the mice M1-M7. Scale bar is 100μm

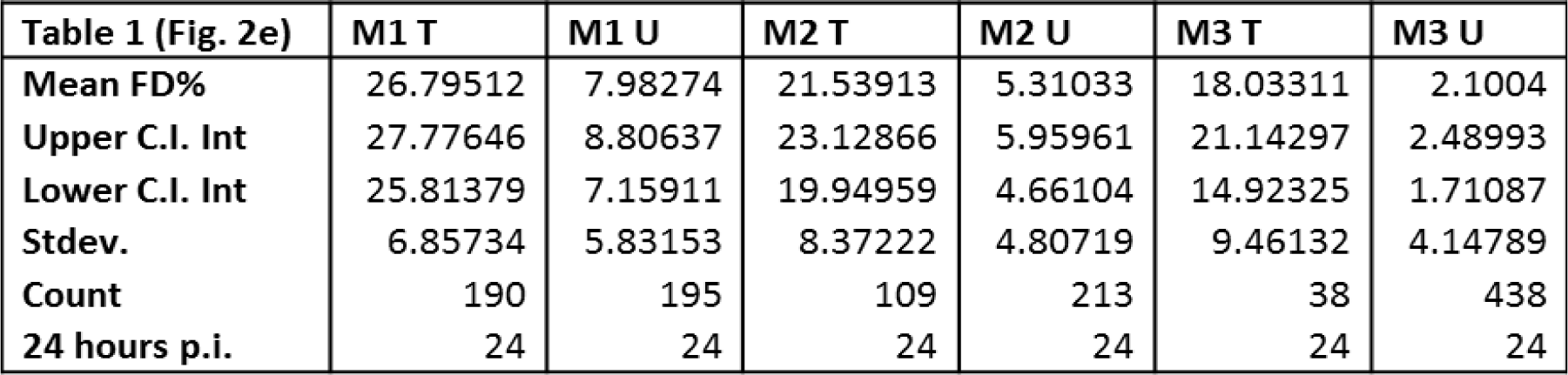

| Table 1 (Fig. 2e) | M1 T | M1 U | M2 T | M2 U | M3 T | M3 U |
| --- | --- | --- | --- | --- | --- | --- |
| Mean FD% | 26.79512 | 7.98274 | 21.53913 | 5.31033 | 18.03311 | 2.1004 |
| Upper C.I. Int | 27.77646 | 8.80637 | 23.12866 | 5.95961 | 21.14297 | 2.48993 |
| Lower C.I. Int | 25.81379 | 7.15911 | 19.94959 | 4.66104 | 14.92325 | 1.71087 |
| Stdev. | 6.85734 | 5.83153 | 8.37222 | 4.80719 | 9.46132 | 4.14789 |
| Count | 190 | 195 | 109 | 213 | 38 | 438 |
| 24 hours p.i. | 24 | 24 | 24 | 24 | 24 | 24 |

| Table 2 (Fig. 3d) | M4 | M5 | M6 | M7 |
| --- | --- | --- | --- | --- |
| Mean FD% | 21.62319 | 8.05126 | 30.10296 | 4.38815 |
| Upper C.I. Int | 22.65215 | 9.10947 | 31.37297 | 5.09102 |
| Lower C.I. Int | 20.59424 | 6.99306 | 28.83296 | 3.68528 |
| Stdev. | 10.44151 | 8.74878 | 10.61917 | 6.19277 |
| Count | 374 | 398 | 265 | 271 |
| 2 hours p.i. | 2 | 2 | 2 | 2 |

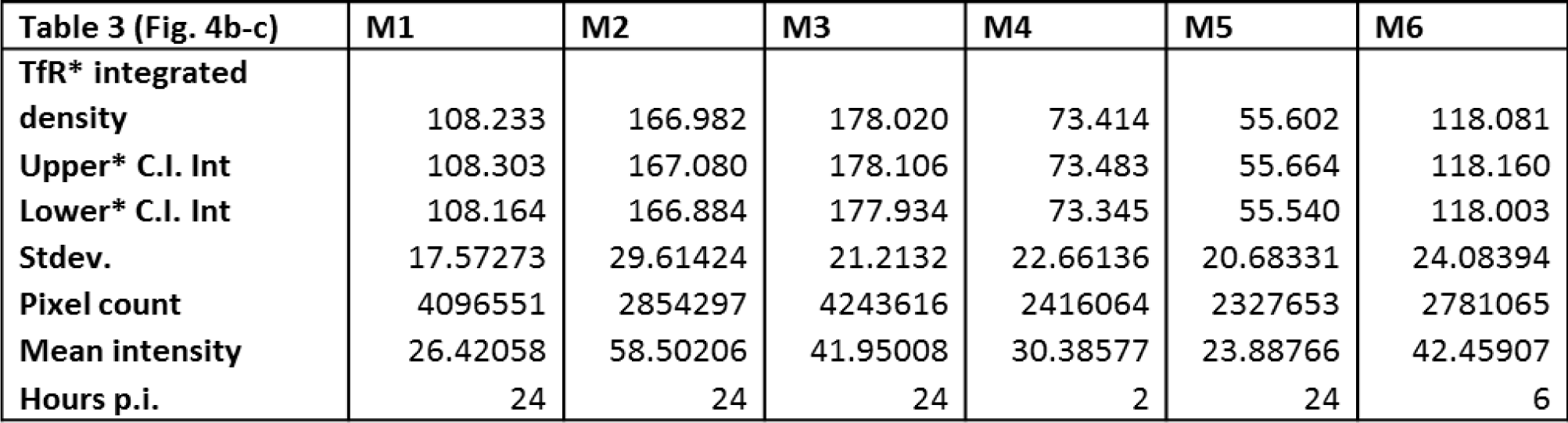

| Table 3 (Fig. 4b-c) | M1 | M2 | M3 | M4 | M5 | M6 |
| --- | --- | --- | --- | --- | --- | --- |
| TfR* integrated density | 108.233 | 166.982 | 178.020 | 73.414 | 55.602 | 118.081 |
| Upper* C.I. Int | 108.303 | 167.080 | 178.106 | 73.483 | 55.664 | 118.160 |
| Lower* C.I. Int | 108.164 | 166.884 | 177.934 | 73.345 | 55.540 | 118.003 |
| Stdev. | 17.57273 | 29.61424 | 21.2132 | 22.66136 | 20.68331 | 24.08394 |
| Pixel count | 4096551 | 2854297 | 4243616 | 2416064 | 2327653 | 2781065 |
| Mean intensity | 26.42058 | 58.50206 | 41.95008 | 30.38577 | 23.88766 | 42.45907 |
| Hours p.i. | 24 | 24 | 24 | 2 | 24 | 6 |

| Table 4 (Fig. 5b-c) | M1 | M2 | M3 | M4 | M5 | M6 |
| --- | --- | --- | --- | --- | --- | --- |
| Tf* integrated density | 50.754 | 19.398 | 11.208 | 27.208 | 53.778 | 82.447 |
| Upper* C.I. Int | 50.783 | 19.414 | 11.223 | 27.241 | 53.804 | 82.479 |
| Lower* C.I. Int | 50.726 | 19.382 | 11.194 | 27.175 | 53.752 | 82.416 |
| Stdev. | 15.88699 | 14.73458 | 17.49876 | 27.60946 | 14.43941 | 13.70937 |
| Pixel count | 833335 | 311563 | 182511 | 373580 | 859653 | 1365320 |
| Mean intensity | 60.90517 | 62.25992 | 61.41118 | 72.83048 | 62.55752 | 60.38671 |
| Hours p.i. | 24 | 24 | 24 | 2 | 24 | 6 |

| Table 5 (Fig 4c & 5c) | M1 | M2 | M3 | M4 | M5 | M6 |
| --- | --- | --- | --- | --- | --- | --- |
| Mean FD% | 26.79512 | 21.53913 | 18.03311 | 21.62319 | 34.38737 | 40.01194 |
| Upper C.I. Int | 27.77646 | 23.12866 | 21.14297 | 22.65215 | 35.77929 | 40.92023 |
| Lower C.I. Int | 25.81379 | 19.94959 | 14.92325 | 20.59424 | 32.99545 | 39.10365 |
| Stdev. | 6.85734 | 8.37222 | 9.46132 | 10.44151 | 8.91474 | 8.1538 |
| Pixel count | 190 | 109 | 38 | 374 | 160 | 312 |
| Hours p.i. | 24 | 24 | 24 | 2 | 24 | 6 |

